# Microfluidic-based high-throughput isolation enhances the recovery of novel strains and diversity from Arctic soil microbiome

**DOI:** 10.64898/2026.06.04.730176

**Authors:** Li Liao, Tingyi Lai, Wanning Jiang, Zedong Duan, Fang Peng, Siqi Zhang, Peiyuan Sun, Yunpeng Zhao

## Abstract

Microbial cultivation remains essential for understanding the physiology, ecology, and biotechnological potential of environmental microbes, yet conventional plate-based methods (CPM) recover only a minute fraction of the environmental microbiome. Polar regions, particularly Arctic soils, represent unique reservoirs of “microbial dark matter” that remain challenging to cultivate, owing to oligotrophic conditions, low temperatures and freeze-thaw cycles that impose severe physiological constraints on microbial growth. Here, we report the first systematic application of microfluidic droplet technology (MDT) to Arctic active-layer soil microbiota and benchmark its performance against CPM using identical starting cell numbers, R2A medium, and incubation at 15°C. MDT achieved 6.5- to 8.1-fold higher recovery rates than CPM and improved isolation throughput by >180-fold. Near-full-length 16S rRNA gene sequencing (PacBio) revealed that MDT recovered significantly higher taxonomic richness across all taxonomic levels, with 256 genera detected in the high-cell-input group (DropAS_H) versus 211 in the corresponding plate group (PlateAS_H). Notably, MDT yielded a more even community distribution, significantly reducing the dominance of fast-growing copiotrophs such as *Pseudomonas* and *Flavobacterium*. Moreover, approximately 50% of sequences from MDT were affiliated with potential novel species (<98.46% identity to type strains), and 27% with potential novel genera (<95% identity). Strain verification by Sanger sequencing confirmed 12 of 17 isolates as candidate novel species, among which one strain represented a potential novel genus within Devosiaceae. This study demonstrates that MDT is a powerful platform for accessing the uncultured majority of polar soil microbiota and establishes a pipeline for high-throughput isolation of novel cold-adapted bacteria.

**IMPORTANCE:** Arctic soils harbor a vast reservoir of microbial diversity that remains largely inaccessible due to the extreme oligotrophic conditions and low temperatures characteristic of polar environments, leading to slow growth rates and extended lag phases in most microbes. Conventional plate-based methods (CPM) inherently favor fast-growing copiotrophs while suppressing rare or slow-growing lineages. Here we demonstrate that microfluidic droplet technology (MDT) overcomes these fundamental constraints, representing its first systematic application to polar microbiology. By physically isolating individual cells into nanoliter-scale bioreactors, MDT mitigates interspecific competition, thereby releasing slow-growing and oligotrophic taxa that are otherwise outcompeted in bulk cultures. The water-in-oil emulsion format further enables extended low-temperature incubation without evaporative loss or airborne fungal contamination, issues that frequently compromise long-term plate-based cultivation of Arctic samples. Relative to CPM, MDT increased recovery rates by >6-fold and isolation throughput by >180-fold, while markedly enhanced both taxonomic richness and evenness. Exclusively recovered by MDT, the oligotrophic genus *Caulobacter* and numerous cold-adapted genera underscore that MDT accesses physiologically distinct fractions of the cryospheric microbiome. Furthermore, the integration of near-full-length 16S rRNA gene sequencing with MDT cultivation assessment provided substantially improved phylogenetic resolution for species-level identification and novel taxon delineation. Collectively, these findings establish MDT as a transformative platform for cryospheric culturomics, accelerating the construction of comprehensive polar strain collections essential for understanding cold-adaptation mechanisms and exploiting the biotechnological potential of Earth’s frozen microbiomes.

## INTRODUCTION

It is widely recognized that <1% of environmental microbes are readily culturable using conventional plate-based methods (CPM), a phenomenon termed the “great plate count anomaly” (1). The remaining uncultured species constitute “microbial dark matter” that awaits discovery (2, 3). Although culturability can exceed 1% under certain circumstances using improved methods (4, 5), the majority of species remain uncultivated (6), especially given the continuously expanding diversity revealed by high-throughput deep sequencing (7). Based on 16S rRNA gene sequencing, approximately 400,000 prokaryotic species comprising about 60,000 genera have been detected (6, 8). Despite this immense diversity, most cultivated bacterial species belong to only four bacterial phyla: Bacteroidota, Pseudomonadota, Bacillota and Actinomyceota (6), highlighting a profound cultivation bias. While advanced single-cell sequencing and multi-omics have expanded our understanding of uncultured microorganisms (9), the limited success of cultivation greatly hinders in-depth investigation of physiology, phenotypes, real ecological functions, and microbial interactions, as well as utilization in industry. Therefore, increased culture efforts combined with culture-independent technologies are required to better understand the uncultured majority.

Culturability varies markedly among environments. For instance, gut microbiomes usually exhibit relatively high cultivation success, with 35 to 65% of molecular species detected by sequencing have representative strains in culture (10). In contrast, extreme environments such as polar regions often show substantially lower success rates. For instance, CPM recovered only 6.37% of bacterial genera detected by metabarcoding in Arctic permafrost soils (11, 12). Polar regions, encompassing the Arctic and Antarctica, are considered unique reservoirs of novel microbial species. Arctic soils, particularly permafrost and active layer soils, are expected to harbor a large pool of psychrophilic and psychrotolerant microorganisms. These microbes are pivotal to climate-relevant biogeochemical cycling (13, 14) and are also promising sources of functional strains and enzymes suited to bioremediation and low-temperature biotechnological applications (15). Accessing the “microbial dark matter” in polar regions is therefore critical for understanding survival mechanisms, metabolic activities, and biogeochemical cycling in the cryosphere, as well as for discovering novel biotechnologically valuable traits.

However, polar regions are characterized by persistently extreme environmental challenges, including low temperatures, prolonged freezing, freeze-thaw cycles, oligotrophy, desiccation, and strong seasonal UV radiation and light regimes (14, 16, 17), all of which impose severe constraints on microbial activity and growth. Such conditions select for cold-adapted, slow-growing microbial populations that often enter dormant states, maintain low metabolic rates, and exhibit extended lag phases, making them particularly difficult to revive and cultivate under standard laboratory conditions and time scales (18, 19). Consequently, advanced cultivation technologies with higher throughput and efficiency are urgently needed for the isolation and cultivation of microbial species from polar regions.

Cultivation efforts in polar microbiology have revealed new taxa and functions, but the accessible diversity remains limited. Isolation and cultivation of microbes from polar environments usually require long incubation periods (up to several months) at low temperatures (20–22). Traditional plate and liquid enrichment cultures tend to favor relatively fast-growing, copiotrophic lineages, while many rare or strictly cold-adapted groups remain uncultured or underrepresented (22, 23). Previous studies have applied CPM to successfully isolate bacteria from Arctic soils (e.g., (11, 14, 24–26)). Meanwhile, other innovative *in situ* cultivation methods including cryo-iplate (26), diffusion chambers (12), iTip and iPore (12) have also been applied to Arctic samples such as lake sediments and soils. These methods recovered different species from plate cultivation but showed limited improvement in cultivation efficiency (26). Despite decades of cultivation efforts, the vast majority of Arctic soil microorganisms remain uncultivated and uncharacterized. Compared with fungi, bacteria showed an even lower rate of culturability in the Arctic soils (11). Therefore, the current gap between the vast uncultured diversity of Arctic soils and the limited cultivation success underscores the need for enhanced methodological throughput.

Microfluidic droplet technology (MDT) has emerged as a transformative platform for microbial cultivation that addresses many limitations of CPM. This technology enables high-throughput encapsulation of single cells into picoliter- to microliter-sized droplets, creating isolated microenvironments that function as miniature bioreactors (27–29). In this way, MDT eliminates interspecific competition by physically isolating individual cells, thereby preventing fast-growing strains from outcompeting slow-growing or dormant cells. Applications of MDT in non-polar soils and gut microbiomes have demonstrated improved cultivation outcomes, including higher richness and recovery of unique genera, compared to CPM (28–30). Moreover, MDT enables the generation of millions of droplets per experiment, significantly increasing the probability of capturing rare and slow-growing taxa. It is particularly suitable for extended incubation period at low temperatures, as the oil emulsion greatly reduces evaporation. Collectively, these characteristics suggest that MDT should be suitable for cultivating recalcitrant Arctic soil microorganisms. Nevertheless, to our knowledge, MDT has not yet been applied to Arctic soil microbial isolation and cultivation.

Given the persistent challenges in cultivating Arctic soil microorganisms and the demonstrated capabilities of MDT in accessing previously uncultivated microbial diversity, this study aims to systematically evaluate the performance of droplet-based single-cell isolation and cultivation for Arctic soil microbiota relative to CPM. Through optimization of encapsulation and incubation protocols tailored to cold-adapted, slow-growing bacteria, we aim to establish a high-efficiency pipeline capable of retrieving taxa refractory to standard plate cultivation. This approach is expected to substantially expand the catalog of cultured polar microorganisms, providing critical strain resources to investigate the Arctic “microbial dark matter” and to discover cold-adapted species with biotechnological potential.

## MATERIALS AND METHODS

### Sampling and soil morphology

During the Chinese Arctic Scientific Research Expedition at the Yellow River Station in September 2023, active-layer soil samples were collected near Ny-Ålesund on Spitsbergen Island, Svalbard (Figure 1). This region experiences seasonal freeze–thaw cycles under the influence of climate change, with a mean annual permafrost temperature of –2.5°C (14). The soils in this area are characterized by oligotrophic conditions with low nutrient availability, typical of high Arctic environments (14). The soil exhibited a coarse, gravelly texture, with abundant coarse sand and gravel particles interspersed throughout a fine mineral matrix. The surface layer was poorly developed, with minimal organic accumulation, and comprised a heterogeneous mixture of angular rock fragments and loose mineral soil. The soil structure was weakly aggregated, consistent with recently exposed glacial till or permafrost active-layer material. At the time of sampling, bryophyte-dominated vegetation was visible in the background, whereas the sampling area itself showed minimal bryophyte colonization. The top 1 cm was removed using a sterile shovel, and soil samples were collected into sterile plastic bags and stored at 4°C until transport to the laboratory in China.

**Figure 1.**
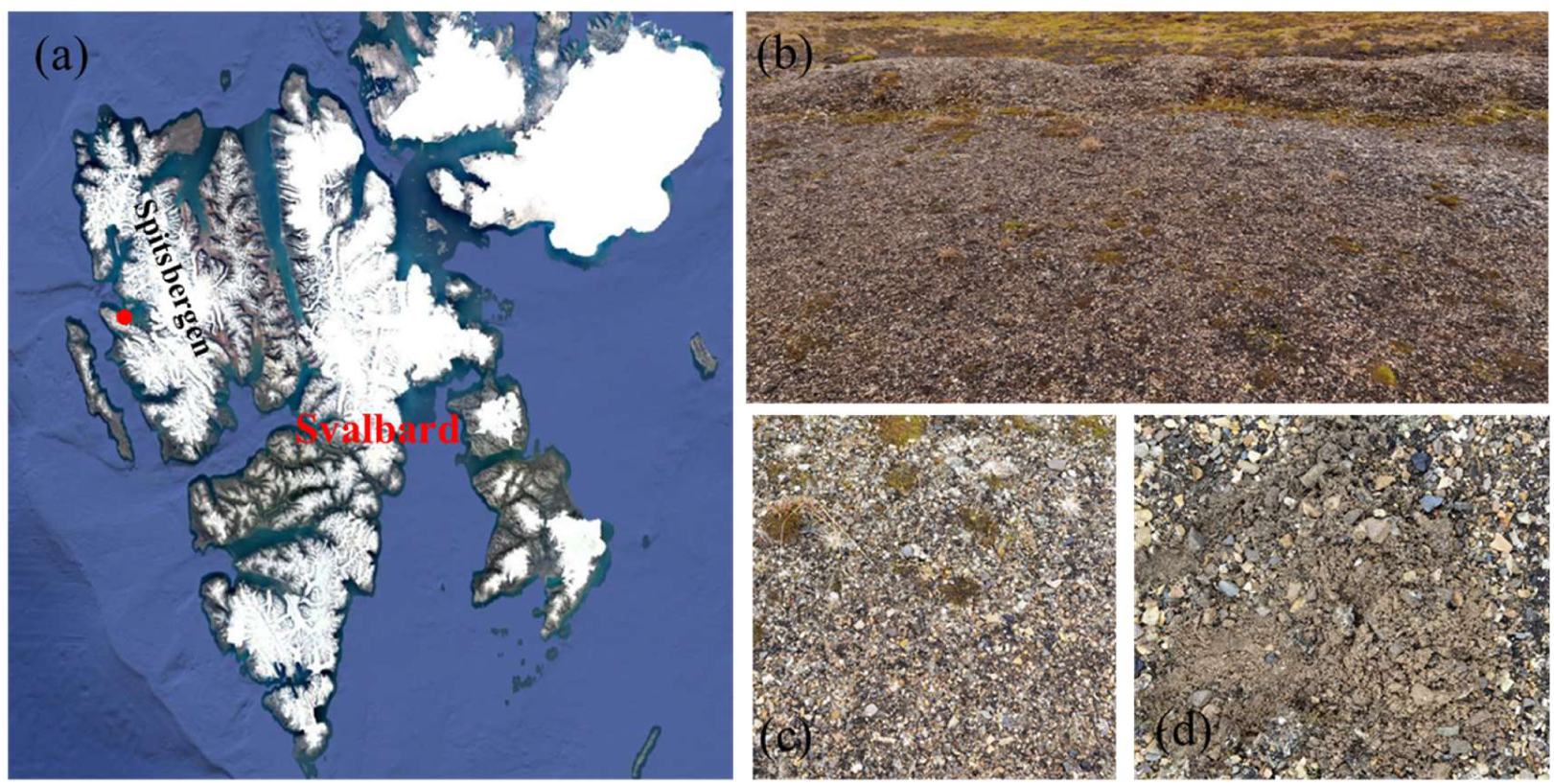
Sampling site information. (a) The sampling site, indicated by a red dot, on Spitsbergen Island in Svalbard, in the Norwegian Arctic. (b) Landscape of the permafrost active layer with sparse vegetation in the background. (c) Close-up view of the soil before sampling. (d) Soil morphology after removal of the top 1 cm layer using a sterile shovel.

### Cell extraction and pre-evaluation using CPM

Approximately 1.5 g of soil sample was weighed and suspended in a sterile tube containing 15 ml filtered normal saline (0.9% NaCl). All the following processes were performed in a cold room at 4℃. The soil suspension was vortex-mixed for 5 minutes and then sonicated for 10 minutes to facilitate cell detachment from soil particles. Following sonication, the suspension was left to settle at 4℃ for approximately 2 hours to allow sedimentation of soil debris. The supernatant was diluted 10-fold before cell counting under a microscope. Final serial dilutions of the supernatant at 1:1, 1:10, 1:100, and 1:1000 were prepared for plate colony-forming unit (CFU) counting. To enable direct comparison between CPM and MDT, we used R2A medium (composition: 0.5 g/L yeast extract, 0.5 g/L proteose peptone, 0.5 g/L casamino acids, 0.5 g/L glucose, 0.5 g/L soluble starch, 0.3 g/L sodium pyruvate, 0.3 g/L K₂HPO₄, 0.05 g/L MgSO₄) (31), a formulation commonly used for microbial isolation from polar samples. Aliquots (100 μl) of the serially diluted suspension were spread onto R2A agar plates and incubated at 15℃ with daily inspection for 30 days. Plates with approximately 50∼100 CFU were used to calculate the viable cell recovery rate, giving approximately 0.04%, substantially lower than the commonly recognized 1%. This value was used as a reference for microdroplet preparation.

### Isolation and cultivation using MDT and CPM

Cell suspension from the above procedure was used for MDT. A total of approximately 4.5 ×10^6^ cells were finally utilized to generate droplets (labeled DropAS_L) following the Poisson distribution formula published previously (32) and the above initial cultivation rate based on CPM. Meanwhile, a 10-fold higher cell input (labeled DropAS_H) was prepared to enhance recovery. Cells were suspended in R2A medium to generate microdroplets using the single-cell microliter-droplet screening system (MISS Cell) at TMAXTREE Biotech (Wuxi, China) (32). The droplet volume was approximately 2.5 nL, and the cell encapsulation followed Poisson distribution principles to maximize single-cell occupancy. A total of 11,895 and 11,534 droplets were generated for DropAS_L and DropAS_H, respectively. All droplets were separated by air and stored in coiled transparent Teflon tubes equipped with the MISS Cell system and incubated at 15℃.

Meanwhile, equivalent cell numbers from DropAS_L and DropAS_H were diluted accordingly and spread on R2A agar plates as controls, labeled PlateAS_L and PlateAS_H, respectively. Both DropAS_L and PlateAS_L, starting with the same number of cells, were incubated at 15℃ for 30 days, representing the lower-cell-input, longer-incubation regimen. DropAS_H and PlateAS_H, containing 10-fold higher cell numbers, were incubated at 15℃ for 17 days, representing the higher-cell-input, shorter-incubation regimen.

### Microdroplet screening and strain picking

At the end of each incubation period, OD_600_ of microdroplets was measured automatically as previously described using the MISS Cell system (32). Microdroplets showing values above the OD_600_ threshold were considered to contain significantly grown cells and were collected into 96-well plates. Up to 100 μl of R2A liquid medium was added to each well to maintain cell viability. The 96-well plates were then stored at 4℃ for further downstream analysis. As to CPM, CFU were counted at the end of incubation for each agar plate, and the plates were then stored at 4℃ for downstream analysis.

### High-throughput 16S rRNA gene sequencing

To rapidly evaluate cultivation results, 10 μl of cell suspension from each well of the 96-well plates was pooled separately for the DropAS_L and DropAS_H regimens. Notably, each well was mixed by pipetting before sampling to avoid cell sedimentation. The pooled cells were centrifuged to remove liquid medium and stored at −80℃ before sequencing. Meanwhile, all colonies grown on agar plates were collected using an inoculation loop with an appropriate amount of sterile 0.9% NaCl solution and transferred to tubes. Liquid was removed by centrifugation before freezing at −80℃.

DNA was extracted using the MagBeads FastDNA Kit for Soil (MP Biomedicals, CA, USA) for all samples. To improve the resolution of species identification, we used 27F (5′–AGAGTTTGATCCTGGCTCAG–3′) and 1492R (5′–GGTTACCTTGTTACGACTT–3′) primers to amplify the near-full-length 16S rRNA gene and sequenced the amplicons using a PacBio Sequel platform at Shanghai Personalbio Technology CO., LTD. (Shanghai, China).

### PacBio sequence analysis

Raw data from PacBio Sequel platform were processed using CCS v4.0.0 software (--min-length 500, --max-length 3000, with default parameters for other parameters) (https://github.com/PacificBiosciences/ccs) (33), generating highly accurate single-molecule consensus reads with base quality values. FASTQ files were demultiplexed based on barcode sequences at both ends of the reads. Barcodes were removed, and reverse-complementary sequences were converted to forward orientation according to the primer sequences.

Near full-length PacBio sequences were processed using the QIIME 2 pipeline (version 2024.2.0) (34). In brief, sequences were imported into QIIME 2 as single-end reads and processed through the following steps: (1) trimming and quality filtering using the cutadapt trim-single plugin with a minimum length threshold of 1,200 bp (--p-minimum-length 1200); (2) dereplication of trimmed sequences using the vsearch dereplicate-sequences plugin to collapse identical sequences; (3) chimera detection and removal using the vsearch uchime-denovo plugin. The resulting high quality, non-chimeric sequences were used for downstream analysis. Taxonomic classification was performed using the feature-classifier classify-sklearn plugin against a SILVA reference database (silva v138.1, full-length 16S rRNA genes).

### Comparison with type strains

To investigate the potential novelty of isolates retrieved from both CPM and MDT, a database of 16S rRNA gene sequences from type strains was constructed. In brief, a list of all available type strains at the time of this work (January 15^th^, 2026) was downloaded from LSPN (List of Prokaryotic names with Standing in Nomenclature) (https://lpsn.dsmz.de/downloads), an authoritative online database providing information on type strains validly published according to the International Code of Nomenclature of Prokaryotes (ICNP) (35). Meanwhile, all 16S rRNA gene sequences were downloaded from the NCBI database. In-house Python scripts were used to extract 16S rRNA sequences according to the LSPN list and to filter matched sequences containing six consecutive N’s to retain high quality for downstream analysis. The resulting type strain reference database (retaining 19,958 sequences between 600 and 1,600 bp) was used for BLASTn comparison. High-quality, non-chimeric near full-length 16S rRNA gene sequences from this study were compared against the type strain database using BLASTn with parameters -evalue 1e-5 and -max_target_seqs 1. The 16S rRNA genes of potential novel taxa were aligned using MAFFT v7.520 (36). Phylogenetic trees were reconstructed using IQ-TREE version 2.2.6 with 1,000 ultrafast bootstrap replicates (-bb 1000) and the best-fitting substitution model determined automatically by the ModelFinder Plus (-m MFP) (37). Phylogenetic trees were annotated using the interactive iTOL online platform (38).

### Strain identification and verification

To verify the sequencing results, microdroplets were randomly selected and further purified on R2A agar plates. Colonies with distinct morphologies were chosen for genomic DNA extraction and 16S rRNA gene amplification using primers 27F and 1492R for Sanger sequencing. Sequences were identified by submitting to the EzBioCloud server (https://www.ezbiocloud.net) (39).

## RESULTS

### MDT achieves higher recovery than CPM

The results demonstrated a dramatic improvement in isolate recovery with MDT. Specifically, the PlateAS_L group (15℃, 30 days) yielded approximately 160 visible CFU by the end of incubation, whereas the PlateAS_H group (15℃, 17 days) produced approximately 450 CFU (Figure 2). Notably, the PlateAS_L group exhibited substantial fungal contamination, likely due to the prolonged 30-day incubation. In contrast, the DropAS_L group (15 °C, 30 days) yielded up to 1,300 microdroplets with visible growth, while the DropAS_H group (15 °C, 17 days) yielded 2,939 microdroplets. Given that both CPM and MDT started with the same number of input cells per comparison, the recovery rate of MDT was calculated to be 8.1-fold (1,300/160) and 6.5-fold (2,939/450) higher than that of CPM, respectively.

**Figure 2.**
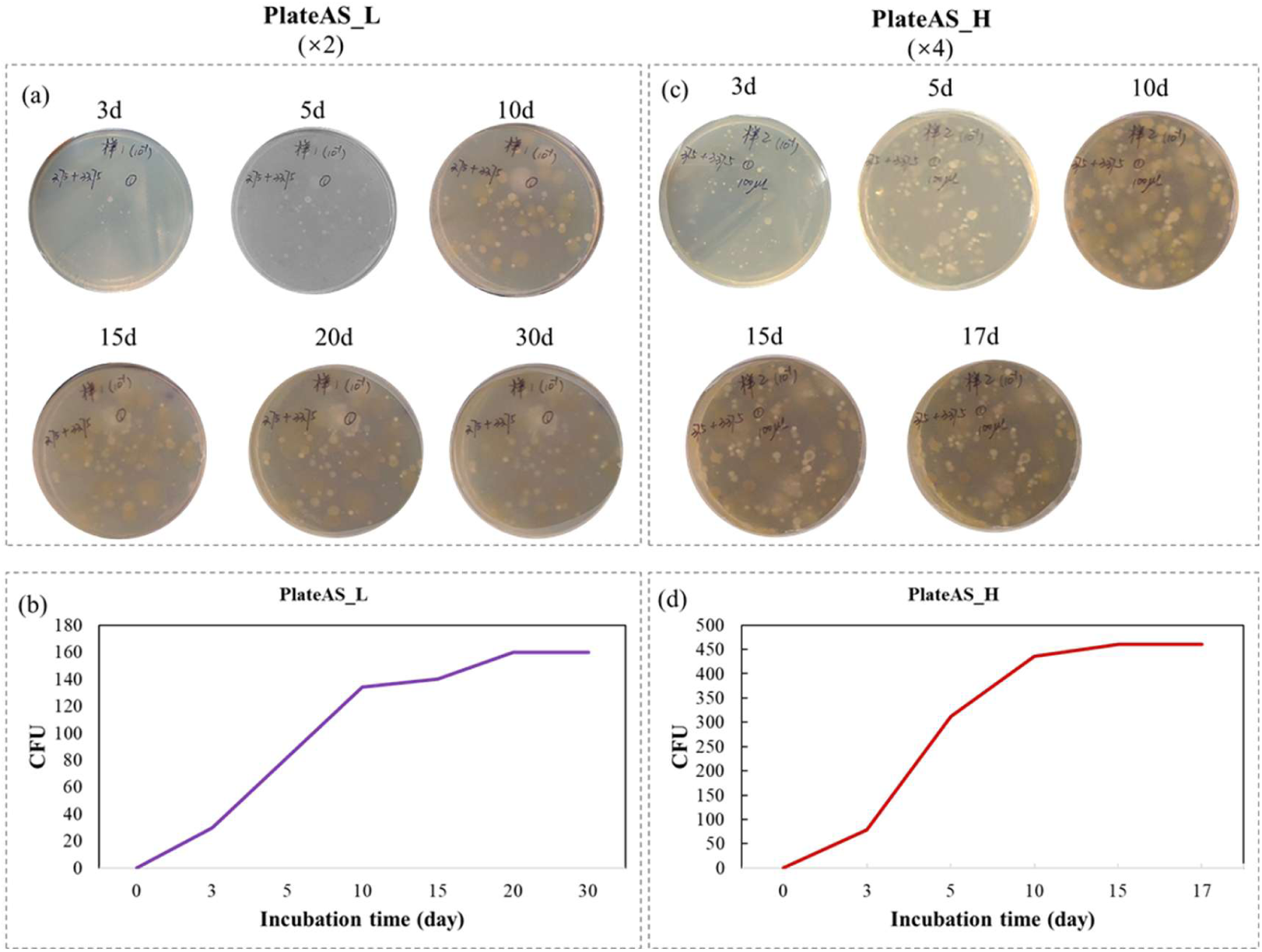
Standard plate culture technique results. (a) Cells equivalent to those used in DropAS_L were equally spread onto two plates; one representative plate is shown. (b) Total CFU on the two plates in (a) were counted manually at various incubation times. (c) Cells equivalent to those used in DropAS_H were equally spread onto four plates; one representative plate is shown. (d) Total CFU on the four plates in (c) were counted manually at various incubation times.

### MDT improves isolation throughput and efficiency

MDT substantially outperformed CPM in both efficiency and throughput. Cell dispersion and isolation were fully automated using the MISS Cell system, operating at a throughput of 5,000–10,000 droplets per hour, which is considerably more efficient than manual spreading and colony picking on agar plates (Figure 3). In this study, following MDT processing of two batches, up to 4,239 microdroplets containing isolates that exhibited significant growth above the threshold designated by the MISS Cell system were collected. Based on average manual workload, isolating an equivalent number of purified strains using CPM would require at least 60 days of work by a single person. In contrast, MDT accomplished the same task in only 8 hours, representing a more than 180-fold improvement in efficiency (Figure 3). Furthermore, MDT consumed substantially fewer consumables, reagents and medium than CPM, and required considerably less space for incubation and other associated processes.

**Figure 3.**
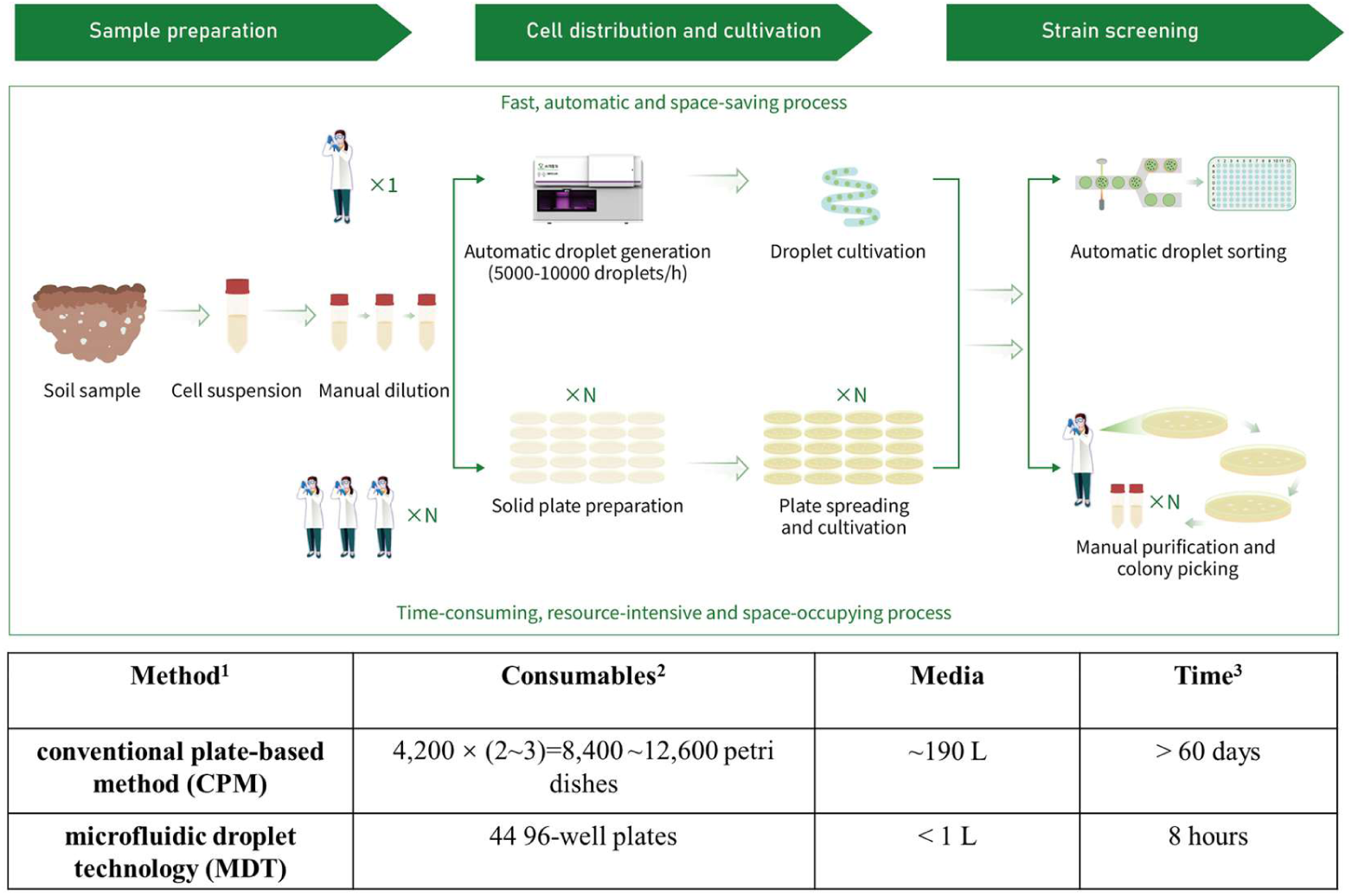
Comparison of two different approaches for isolation, cultivation and screening: conventional plate-based method (CPM) and high throughput microfluidic droplet technology (MDT). ^1^The calculation is based on the recovery numbers of colonies from plates and of microdroplets achieved in this study, assuming a target of obtaining 4,200 strains using either method. ^2^The standard plate culture technique typically requires two to three rounds of streak purification to obtain a pure culture. Only the main consumables are included for comparison. ^3^Based on a single person completing 200 plate spreading/streaking operations per day, the minimum time required for plate preparation and subsequent spreading/streaking was calculated.

### MDT recovers higher taxonomic richness and evenness at all levels

To enable comparison of species composition recovered by MDT and CPM at higher phylogenetic resolution, over 70,000 near-full-length 16S rRNA gene sequences were generated per experiment using the PacBio Sequel platform. After filtering, trimming, dereplication, and chimera removal using the QIIME 2 pipeline, 23,423–39,899 final ASVs longer than 1,200 bp were retained for downstream analysis (Table 1). Within both MDT and CPM, high-cell-input groups (DropAS_H and PlateAS_H) yielded a greater number of ASVs than corresponding low-cell-input groups (DropAS_L and PlateAS_L). Sequences were annotated at various taxonomic levels from phylum to genus (Table 1). Across all samples, a total of 22 phyla and 283 genera were identified after excluding unclassified taxa. Pseudomonadota, Bacteroidota, Actinomyceota and Bacillota were the dominant phyla. At all taxonomic levels, DropAS_H always ranked first among them. Overall, MDT groups contained a higher number of taxa than CPM controls across all taxonomic levels. Specifically, DropAS_H contained 256 genera (including unclassified genera), 45 more than the 211 detected in PlateAS_H. Similarly, DropAS_L yielded 59 more genera than PlateAS_L. At the phylum level, 3 and 2 additional phyla were detected in DropAS_H and DropAS_L, respectively, compared to PlateAS_H and PlateAS_L.

**Table 1.**
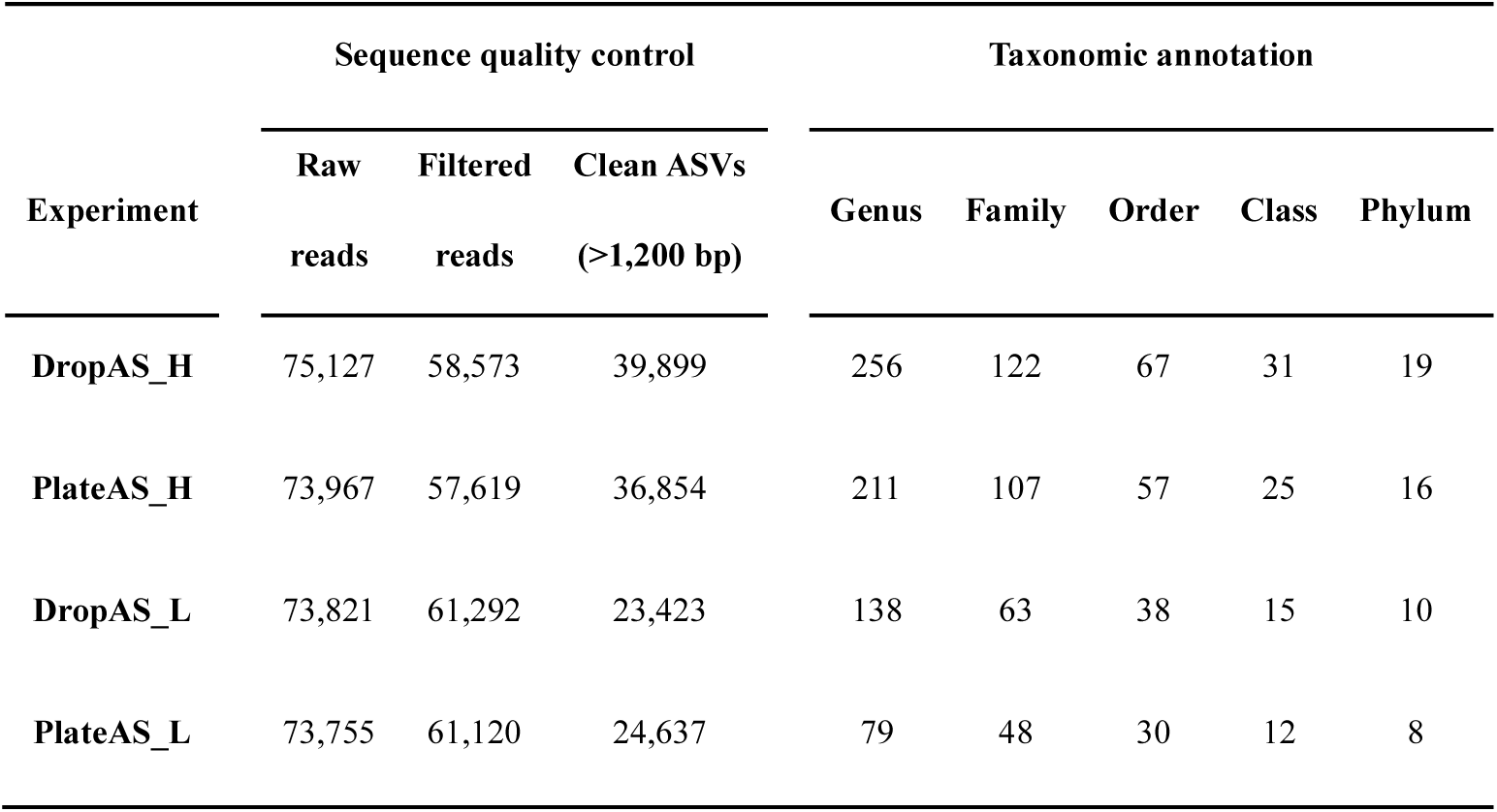
Sequence quality control and taxonomic annotation.

### MDT reduces copiotroph dominance

To focus on the comparison between MDT and CPM, genera recovered across all four gourps were analyzed in detail. The number of genera detected varied considerably, with only 41 genera (11%) shared among the four groups (Figure 4a). The highest number of unique genera was detected in DropAS_H (61, 17%), whereas the lowest was in PlateAS_L (5, 1%), representing a 12.2-fold increase. The second highest number of unique genera was detected in PlateAS_H (56, 15%), followed by DropAS_L (39, 11%).

**Figure 4.**
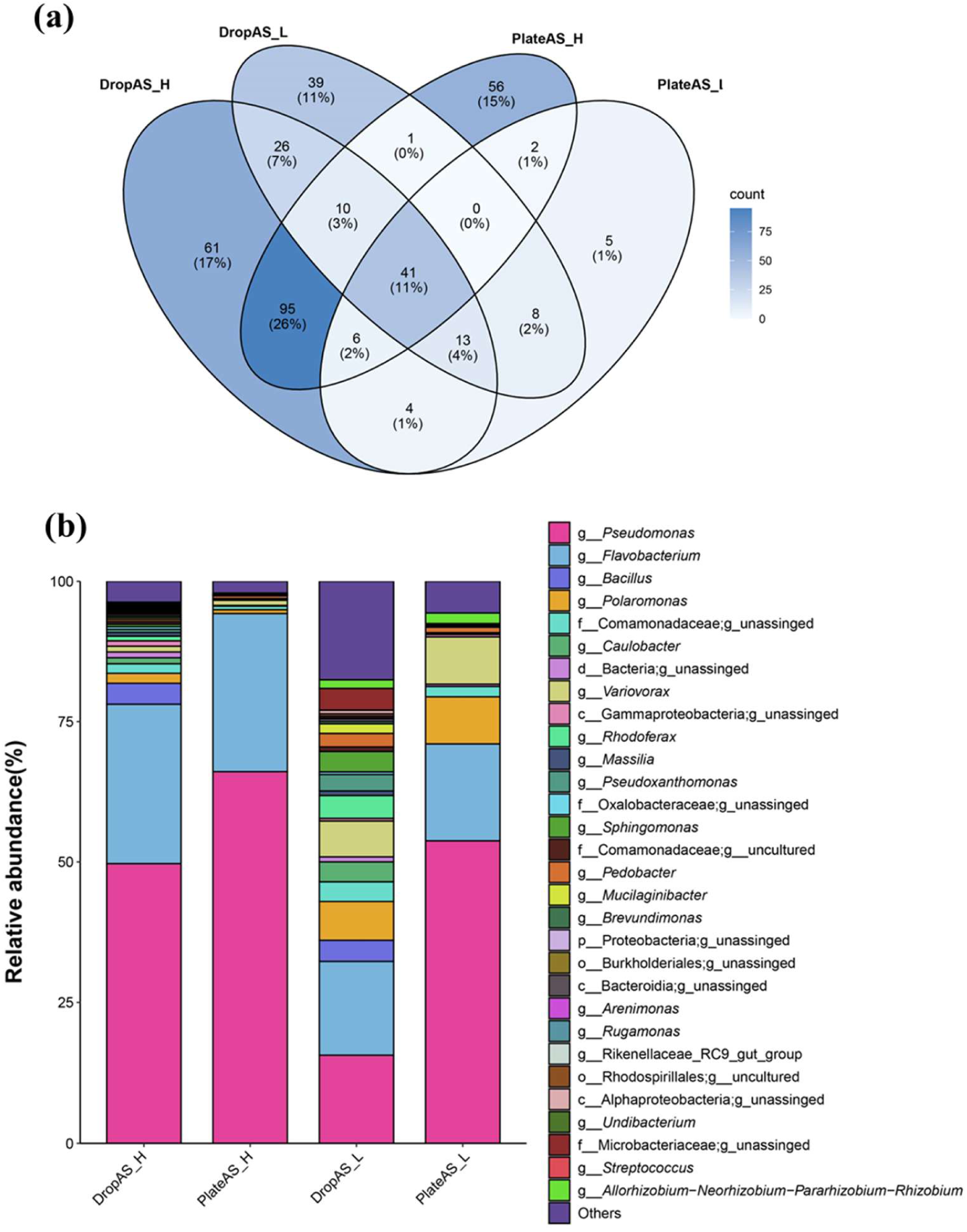
Comparison of genera detected across isolation experiments. (a) Venn diagram of shared genera. (b) Composition and distribution of top 30 genera.

Although two genera, *Pseudomonas* and *Flavobacterium*, dominated in all four experiments, their relative abundances varied markedly (Figure 4b). Notably, *Pseudomonas* and *Flavobacterium* accounted for up to 66.1% and 28.2% of reads in PlateAS_H (the highest), but only 15.6% and 16.6% in DropAS_L (the lowest), respectively. Another apparent difference between MDT and CPM groups was the evenness of genus distribution. MDT groups exhibited both higher genus richness and greater evenness than corresponding CPM controls. Moreover, DropAS_L contained the highest percentage of rare genera (i.e., those outside the top 30 shown in Figure 4b) (17.5% in DropAS_L vs. 2.1% in PlateAS_H).

### MDT and CPM enrich different groups

Different taxa were enriched between MDT (DropAS_L and DropAS_H) and CPM groups (PlateAS_L and PlateAS_H) (Figure 5). Specifically, unclassified bacteria at various taxonomic levels, from domain to family, including Bacteriodia, Pseudomonadota, Oxalobacteraceae, Xanthomonadaceae and Sphingobacteriacaea were much more abundant in MDT (up to >40-fold higher) than in CPM groups. Besides, genera *Bacillus*, *Massllia*, *Mucilaginibacter*, *Brevundimonas*, and *Arenimonas* were most abundant in MDT group (up to >1,600 fold higher). Moreover, genera with relative abundances >0.01%, including *Caulobacter* (1.1%, 3.6%), unclassified Caulobacteraceae (0.05, 0.09%), *Sphingomonas* (0.5%, 3.6%), *Arenimonas* (0.2%, 0.3%), *Cryobacterium* (0.06%, 0.06%), *Pelomonas* (0.06%, 0.3%), *Nocardioides* (0.04%, 0.9%), *Paucibacter* (0.03%, 0.7%), *Sphingopyxis* (0.03%, 0.1%), *Rhizorhapis* (0.03%, 0.3%), *Uliginosibacterium* (0.02%, 0.03%), *Sphingobium* (0.02%, 0.04%), and unclassified Cellulomonadaceae (0.01%, 0.02%), were exclusively detected in the MDT (values in parentheses represent relative abundances in DropAS_H and DropAS_L, respectively) but absent in CPM.

**Figure 5.**
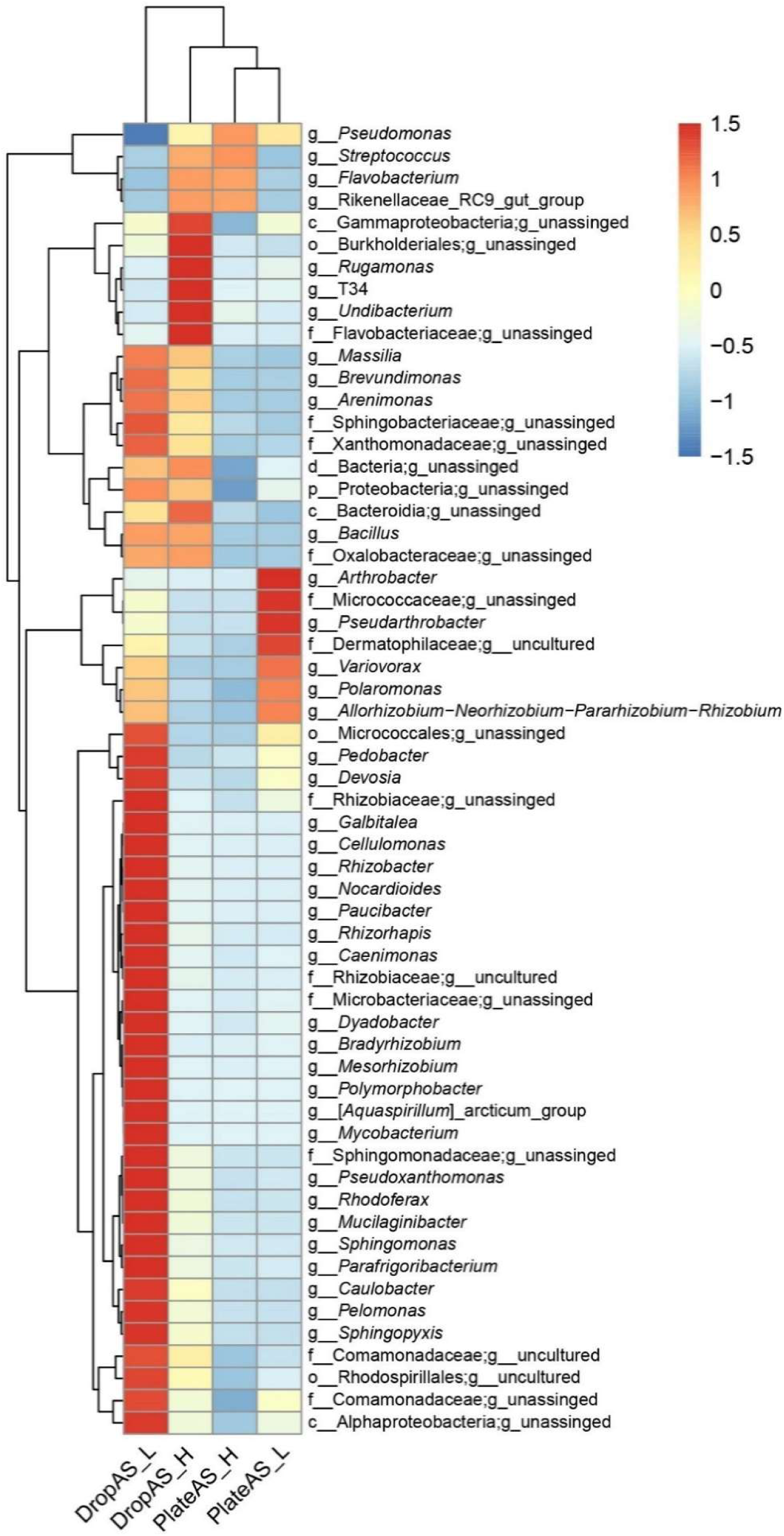
Heatmap of genera with abundance over 0.1%.

Specific taxa were enriched in DropAS_L, including genera *Pedobacter*, *Devosia*, *Galbitalea*, *Cellulomonas*, *Rhizobacter*, *Nocardioides*, *Paucibacter*, *Rhizorhapis*, *Caenimonas*, *Dyadobacter*, *Bradyrhizobium*, *Mesorhizobium*, *Polymorphobacter*, *Mycobacterium*, *Pseudoxanthomonas*, *Rhodoferax*, *Mucilaginibacter*, *Sphingomonas*, *Parafrigoribacterium*, *Caulobacter*, *Pelomonas* and *Sphingopyxis*, as well as unclassified groups from Alphaproteobacteria, Micrococcales, Rhizobiaceae, Microbacteriaceae, Sphingomonadaceae, Comamonadaceae, and Rhodospirillales (Figure 5).

Although the most apparent difference existed between MDT and CPM groups, long incubation tended to enrich specific groups in both DropAS_L and PlateAS_L, including *Pseudarthrobacter*, *Allorhizobium-Neorhizobium -Pararhizobium Rhizobium*, *Polaromonas*, *Variovorax*. In particular, genera *Arthrobacter*, *Pseudarthrobacter*, and unassigned genera in Micrococcaceae and Dermatophilaceae were highly enriched in PlateAS_L.

### MDT recovers more potential novel taxa

To identify potential novel species and genera recovered in the four experiments, 16S rRNA gene sequences were retrieved for validly published type strains available in the LSPN database and used as the database for BLASTn (see Methods). According to the widely used 98.46% and 95% 16S rRNA identity thresholds for novel species and genera (40), 49.5% and 27.2% of sequences that matched type strains in DropAS_H represented respectively potential novel species and genera, which were the highest among all experiments (Table 2). In contrast, the lowest percentages of potential novel genera (22.2%) and species (9.8%) were detected in PlateAS_H. Overall, about 1.6- to 2.2-fold more potential novel species and genera were detected in MDT groups than in CPM groups.

**Table 2.** Statistics of potential novel taxa.

|  | Number of ASVs belonging to potential novel taxa (based on 16S identity) |  |  |  |  |
| --- | --- | --- | --- | --- | --- |
|  | Clean ASVs | Species (< 98.64%) | Genus (< 95%) | Family (< 86.5%) | Order (< 82%) |
| DropAS_H | 39,899 | 19,756 (49.5%) | 10,864 (27.2%) | 436 (1.1%) | 27 (0.07%) |
| PlateAS_H | 36,854 | 8,197 (22.2%) | 3,615 (9.8%) | 246 (0.67%) | 30 (0.08%) |
| DropAS_L | 23,423 | 11,181 (47.7%) | 3,532 (15.1%) | 103 (0.44%) | 1 (0.004%) |
| PlateAS_L | 24,637 | 7,271 (29.5%) | 2,953 (12.0%) | 36 (0.15%) | 0 (0%) |

To further investigate the novelty of sequences recovered from this study, sequences sharing less than 82% identity with type strains, representing potential novel order (40), were pooled to construct a phylogenetic tree with the closest type strains (Figure 6). These included 27 sequences from DropAS_H (belonging to six phyla: Bacillota, Bacteroidota, Pseudomonadota, Kiritimatiellota, Cyanobacteriota, and Verrucomicrobiota), 30 from PlateAS_H (belonging to five phyla: Bacillota, Bacteroidota, Pseudomonadota, Spirochaetota, and Actinomyceota), and one from DropAS_L belonging to Pseudomonadota. More than half of the sequences recovered from this study clustered separately from type strains. In particular, 10 potential clusters were identified as putative novel orders not closely related to known groups, most of which belonged to Bacteroidota. In addition to these 10 potential novel clusters, other sequences from this study may represent novel taxa above the family level from rare phyla such as Spirochaetota, Bacillota, Kiritimatiellota and Verrucomicrobiota.

**Figure 6.**
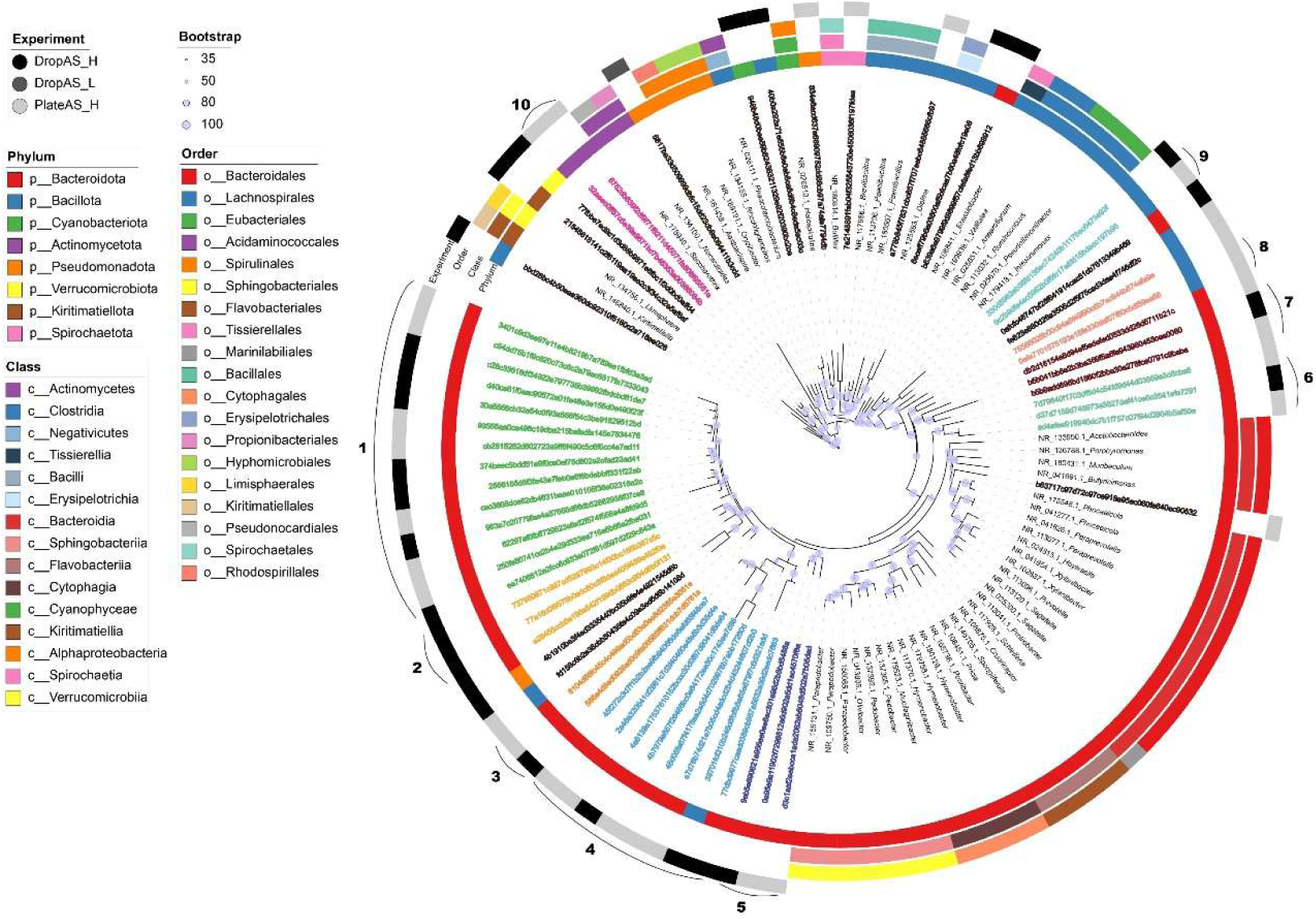
Phylogenetic tree of sequences sharing less than 82% identity with type strains. Four concentric rings (from inner to outer) denote phylum, class, order, and experimental groupings, respectively. Sequences from this study are identified by their Feature IDs, while type strains are labeled with accession numbers and genus names. Leaf node colors reflect different putative clusters, which correspond to the numeric labels and curved lines above the outermost ring (marked “Experiment”). Bootstrap support values are represented as circles on branches, with the size scale explained in the legend on the left. The color key is also provided on the left.

### Strain isolation verification

Fifty microdroplets with varying OD_600_ values, representative of different growth rates, were randomly selected for validation, given the labor-intensive nature of downstream processing and the need for preliminary confirmation of cultivation success. Generally, uniform colonies appeared on R2A plates derived from individual microdroplets, indicating that each droplet likely contained a single cultivated species. A total of 20 colonies with distinct morphologies were chosen for Sanger sequencing. Finally, 17 strains were successfully sequenced, whereas three failed during DNA extraction and sequencing (Table 3). These 17 strains showed 16S rRNA gene sequence identities ranging from 92.92% to 100% with their closest type strains. Among them, 12 out of 17 showed identities below 98.64%, representing potential novel species within *Arthrobacter*, *Bacillus*, *Flavobacterium*, *Mesorhizobium*, *Nocardioides*, *Polaromonas*, and *Sphingomonas*. One strain, labeled as S-114-7H, showed <95% identity with the closest type strain *Devosia insulae* DS-56, standing for a potential novel genus with Devosiaceae of Pseudomonadota. All genera obtained here were also detected in the high-throughput PacBio sequencing results, with C*ellulomonas*, *Devosia*, *Nocardioides*, *Mesorhizobium*, *Sphingomonas*, *Polaromonas*, and *Bacillus* enriched in MDT experiments (Figure 6). These results confirm the reliability of MDT sequencing data and demonstrate the feasibility of MDT for obtaining novel species from Arctic soil samples.

**Table 3.** Identification of strains purified from microdroplets.

| Strain | Closest type strain | Identity | Coverage | Phylum |
| --- | --- | --- | --- | --- |
| S-133-4B | *Cellulomonas humilata* ATCC 25174 | 99.31% | 60.60% | Actinomycetota |
| S-132-11H | *Mesorhizobium norvegicum* 10.2.2 | 97.86% | 99.00% | Pseudomonadota |
|  | *Mesorhizobium loti* NBRC 14779 |  |  |  |
| S-132-10C | *Nocardioides allogilvus* CFH 30205 | 98.37% | 55.40% | Actinomycetota |
| S-114-7H | *Devosia insulae* DS-56 | 92.92% | 98.00% | Pseudomonadota |
| S-114-7G | *Sphingomonas silvisoli* RP18 | 96.27% | 98.00% | Pseudomonadota |
| S-114-7F | *Pelomonas dachongensis* DC23W | 99.06% | 96.00% | Pseudomonadota |
| S-114-7E | *Mesorhizobium norvegicum* 10.2.2 | 99.00% | 98.00% | Pseudomonadota |
| S-112-9D | *Mesorhizobium norvegicum* 10.2.2 | 97.14% | 97.00% | Pseudomonadota |
|  | *Mesorhizobium loti* NBRC 14779 |  |  |  |
| S-112-1B | *Polaromonas jejuensis* NBRC 106434 | 95.74% | 98.00% | Pseudomonadota |
| S-101-1G | *Polaromonas ginsengisoli* Gsoil 115 | 97.19% | 99.00% | Pseudomonadota |
| F-143-8A | *Pseudomonas yamanorum* 8H1 | 100% | 54.60% | Pseudomonadota |
| F-143-3D | *Flavobacterium collinsii* 983-08 | 96.70% | 98.00% | Bacteroidota |
| F-141-G11 | *Arthrobacter ginsengisoli* DCY81 | 96.85% | 98.00% | Actinomycetota |
| F-141-B9 | *Bacillus cereus* ATCC 14579 | 97.73% | 99.00% | Bacillota |
| F-141-A9 | *Bacillus cereus* ATCC 14579 | 98.17% | 98.00% | Bacillota |
| F-114-12D | *Polaromonas ginsengisoli* Gsoil 115 | 98.52% | 98.00% | Pseudomonadota |
| F-113-G9 | *Bacillus cereus* ATCC 14579 | 98.02% | 97.00% | Bacillota |

## DISCUSSION

This study represents the first application and systematic evaluation of MDT for isolation and cultivation of microorganisms from Arctic active-layer soils. Our results demonstrate that MDT not only increased isolation throughput by more than 180-fold compared to CPM, but more importantly, substantially improved the representativeness of recovered community composition. Under identical starting cell numbers and R2A medium conditions, MDT recovered 21–75% more genera than CPM, with a significantly more even community distribution. For instance, the combined relative abundance of dominant copiotrophic genera *Pseudomonas* and *Flavobacterium* decreased from 94.3% in CPM to 32.2% in MDT. These findings align with previous MDT applications in gut microbiomes and non-extreme soils, where droplet confinement effectively alleviates interspecific competition and releases slow-growing or rare taxa that are otherwise outcompeted by fast-growing strains (29, 41). Consequently, high throughput methods like MDT can effectively address the limitations of CPM, particularly for extreme environment samples that harbor slow-growing microbes and require prolonged incubation. Beyond these performance gains, the scalability and efficiency of MDT offer distinct advantages for large-scale investigations. As a powerful complement to meta-omics approaches, it enables the high-throughput construction of microbial strain collections from polar regions. As demonstrated in this study and elsewhere (27, 42), MDT dramatically reduces labor input, increases throughput, and conserves reagents and consumables, thereby providing a practical and robust pipeline for accessing the untapped cultural diversity of Earth’s cryosphere.

Among the MDT-enriched genera, *Caulobacter* is particularly noteworthy. This genus is a classic model for studying asymmetric cell division and polar development, characterized by a distinctive stalk with a holdfast adhesin at its distal end that attatches to solid-liquid interfaces (43). Although *Caulobacter* has long been regarded primarily as an aquatic oligotroph, recent metagenomic analyses challenge this view, demonstrating that its relative abundance in soils is on average fourfold higher than in aquatic environments (44). The successful recovery of *Caulobacter* from Arctic soil in this study supports this revised distribution pattern and further suggests its active adaptation to low-nutrient conditions and periodic freeze-thaw cycles at solid-liquid interfaces. Interestingly, *Caulobacter* was specifically enriched in MDT (1.1% in DropAS_H and 3.6% in DropAS_L) yet completely absent in CPM. One plausible explanation relates to the contrasting physical environments: CPM provides only a solid agar surface, whereas MDT generates a liquid–solid–air interface. This hypothesis is supported by a previous report that *Caulobacter* also colonizes the air–liquid interface (45). Further investigation is needed to fully elucidate the basis for this differential recovery. Nevertheless, this differential recovery underscores the superior capacity of MDT over CPM in accessing specific oligotrophic-adapted bacterial groups from complex environmental samples.

Beyond *Caulobacter*, several genera enriched or exclusively detected in MDT possess clear biotechnological applications and value in studying adaptation to Arctic soil environment. For instances, the genus *Bacillus*, which was entirely absent in PlateAS_H but reached 3.7% in MDT, serves as a major chassis (in particular *B. subtilis*) for industrial protein expression and as plant growth-promoting microbial fertilizers (46). Cold-adapted *Bacillus* strains from Arctic soils could be exploited as platforms for low-temperature enzyme expression or as biofertilizers for cold-region agriculture. The genus *Massilia* was enriched over 1,600-fold in MDT relative to CPM. Multiple novel *Massilia* species have been described from Antarctic streams, lakes, and regoliths, demonstrating strong adaptation to extreme cold and oligotrophic conditions (47, 48). Genera such as *Cryobacterium*, *Mucilaginibacter*, and *Arenimonas* are well-known cold-adapted bacteria widely distributed in cryoconite, glacial ice, and permafrost soils, representing characteristic members of indigenous cryospheric microbial communities (49). Additionally, *Paucibacter* (known for microcystin and nodularin detoxification) (50) and *Sphingopyxis* (capable of pesticide degradation) (51) were enriched in MDT, suggesting potential applications in cryosphere bioremediation. Genera such as *Nocardioides*, *Massilia*, and *Mucilaginibacter* are also recognized for active secondary metabolite production, with potential biomedical value (52). Altogether, the microbial cultures obtained in this study especially from MDT showed great potential in understanding mechanisms behind adaptation to cold and nutrient depletion, and also in future applications in various field.

Different from previous MDT studies, our study applied several different strategies. First, we used a larger volume (∼2.5 nl/droplet) rather than picoliter volumes commonly employed in previous work. Although this reduces the total number of droplets generated, it substantially facilitates downstream processing, including long-term incubation stability, single-droplet dispensing into microwells, and preservation of droplets for subsequent validation and strain purification. This volume trade-off was essential for slow-growing polar microorganisms that require extended incubation. Second, we utilized near-full-length 16S rRNA gene sequencing via PacBio for the first time in MDT cultivation assessment. This approach provides substantially greater phylogenetic resolution and higher confidence in species identification. Previous MDT studies evaluating cultivation outcomes relied almost exclusively on short-read sequencing of partial 16S rRNA gene fragments (∼250–300 bp, typically the V4 region) (e.g., (28, 29)), which provides limited resolution for species-level identification and novel taxon delineation. Third, our study is the first to evaluate MDT applicability for extreme environmental samples from the cryosphere. Different from prior MDT studies conducted under 30∼37°C with short incubation (typically 1-5 days) (29, 41), we optimized the cultivation temperature to 15°C and extended the incubation period up to 30 days. This regimen was necessary to accommodate the slow growth rates and extended lag phases characteristic of cold-adapted soil microbiota. The water-in-oil emulsion system, combined with sealed Teflon tubing incubation, effectively prevented both evaporation and airborne contamination, issues that frequently compromise long-term incubation using CPM. Thus, our results demonstrate that MDT is particularly suitable for environmental samples from the cryosphere, offering a robust platform for long-term, low-temperature incubation without the desiccation and fungal contamination problems commonly encountered in CPM.

Despite the demonstrated advantages of MDT, this study has several limitations that point to future improvements. First, we only used one cultivation medium R2A in this study. Although R2A medium is a standard oligotrophic formulation widely used for microbial isolation from the polar regions (53, 54) and was appropriately chosen here to evaluate the baseline applicability of MDT in Arctic soils, its use alone inevitably excluded chemolithoautotrhophs, strict anaerobes, and taxa requiring specific substrates for growth. Media supplemented by soil extract was shown to increase the cultivation diversity in a previous MDT study (28). Future efforts should therefore expand the cultivation repertoire to include multiple media, as well as variations in oxygen levels and temperatures to capture the full physiological diversity of the cryosphere microbiome, as proposed previously (12). Another constraint lies in the downstream recovery of novel taxa. Although PacBio sequencing revealed substantial proportions of potential novel species (49.5% in DropAS_H) and novel genera (27.2%), the transition from sequencing-based detection to physical isolation of higher-order novel taxa (e.g., novel families or orders) remains challenging. The targetability of downstream analysis is currently limited due to the need to screen thousands of droplets to locate a target strain present as a low-abundance sequence in the pooled library. Future integration of in-droplet identification technologies could bridge this gap by enabling targeted recovery of specific taxa directly from the droplet pool.

Furthermore, as a common challenge faced by MDT in strain isolation, by physically isolating individual cells into droplets, MDT simultaneously disrupts in situ microbial interaction networks, including syntrophy, cross-feeding, and quorum sensing. Thus, the isolation effect of MDT is a double-edged sword: on one hand, it releases slow-growing taxa suppressed by competition; on the other hand, it may prevent the growth of taxa strictly dependent on metabolic interactions. Nevertheless, under the oligotrophic conditions commonly observed in polar environments, competition rather than metabolic cooperation could be the primary force structuring microbial communities (55–57). Theory and empirical evidence from diverse oligotrophic systems indicate that when nutrient availability is chronically low, k-selected oligotrophs consistently outcompete r-selected copiotrophs for scarce resources by maintaining high-affinity uptake systems and low metabolic maintenance costs (55, 56). Consequently, the competitive release afforded by droplet confinement outweighs the loss of potential community interactions, which explains the markedly improved recovery of diversity, quantity, and novelty observed in this study. However, the higher cell input group (DropAS_H) showed higher taxonomic novelty than the lower input group (DropAS_L), suggesting that multi-cell encapsulation at moderate frequency may occasionally capture synergistic micro-colonies or metabolically interdependent cells that would be excluded under strict single-cell encapsulation. This observation hints at a promising future direction: MDT could be adapted for controlled co-cultivation by deliberately increasing multi-cell encapsulation rates (higher Poisson λ), thereby preserving beneficial metabolic interactions while still mitigating the overwhelming competitive suppression seen in bulk plate cultures.

## CONCLUSIONS

This study provides the first systematic evaluation of MDT for the isolation and cultivation of microorganisms from Arctic active-layer soils. Our results demonstrate that MDT substantially outperforms CPM across multiple metrics: recovery rates were 6.5- to 8.1-fold higher, isolation throughput improved by >180-fold, and the recovered community exhibited significantly greater taxonomic richness (256 versus 211 genera) and evenness. The near-full-length 16S rRNA gene sequencing approach enabled robust taxonomic assignment, revealing that approximately 50% of MDT-recovered sequences represented potential novel species and 27% potential novel genera. The differential enrichment of *Caulobacter* and other oligotrophic-adapted genera in MDT but not in CPM highlights that droplet confinement effectively mitigates the competitive dominance of fast-growing copiotrophs, thereby releasing slow-growing and rare taxa that are refractory to standard cultivation.

In future research, expanding the repertoire of cultivation media and incubation conditions will be essential to fully exploit the advantages of MDT and further enhance recovery efficiency and taxonomic coverage. Concurrently, addressing the downstream bottlenecks of strain dereplication and targeted sorting from thousands of droplets remains a critical challenge. We anticipate that integrating MDT with culturomics and multi-omics will accelerate the construction of comprehensive polar strain collections, advancing both fundamental understanding of microbial life under extreme conditions and the biotechnological exploitation of cryospheric resources.

## ACKNOWLEDGEMENTS

The project was supported by the National Key Research and Development Program of China (Grant No. 2022YFC2807501), and the National Natural Science Foundation of China (Grant No. 42476264 and 41976224).

